# Reconciling Conformational Heterogeneity and Substrate Recognition in Cytochrome P450

**DOI:** 10.1101/2020.06.08.139790

**Authors:** B. Dandekar, N. Ahalawat, J. Mondal

## Abstract

Cytochrome P450, the ubiquitous metalloenzyme involved in detoxification of foreign components, has remained one of the most popular systems for substrate-recognition process. However, despite being known for its high substrate specificity, the mechanistic basis of substrate-binding by archetypal system cytochrome P450cam has remained at odds with the contrasting reports of multiple diverse crystallographic structures of its substrate-free form. Here we address this issue by elucidating the probability of mutual dynamical transition to the other crystallographic pose of cytochrome P450cam and vice versa via unbiased all-atom computer simulation. A robust Markov state model (MSM), constructed using adaptively sampled 84 microsecond-long Molecular dynamics simulation trajectories, maps the broad and heterogenous P450cam conformational landscape into five key sub-states. In particular, the MSM identifies an intermediate-assisted dynamic equilibrium between a pair of conformations of P450cam, in which the substrate-recognition sites remain ‘closed’ and ‘open’ respectively. However, the estimate of a significantly high stationary population of closed conformation, coupled with faster rate of open → closed transition than its reverse process, dictates that the net conformational equilibrium would be swayed in favour of ‘closed’ conformation. Together, the investigation quantitatively infers that while a potential substrate of cytochrome P450cam would in principle explore a diverse array of conformations of substrate-free protein, it would mostly encounter a ‘closed’ or solvent-occluded conformation and hence would follow an induced-fit based recognition process. Overall, the work reconciles multiple precedent crystallographic, spectroscopic investigations and establishes how a statistical elucidation of conformational heterogeneity in protein would provide crucial insights in the mechanism of potential substrate-recognition process.

**STATEMENT OF SIGNIFICANCE:** Conformational heterogeneity plays an important role in defining the structural and functional dynamics of the enzymes. While the static three-dimensional crystallographic structures of enzymes solved in different conditions and/or environments are crucial to provide the conformational sub-states of enzymes, these are not sufficient to understand the kinetics and thermodynamics of these sub-states and their role in substrate recognition process. Cytochrome P450cam, the archtypal metalloenzyme, presents such a complex scenario due to prevalent reports of contrasting crystallographic structures of its substrate-free form. This work quantifies the conformational heterogeneity of substrate-free P450cam by exploring the possibility of mutual transition among the crystallographic poses at an atomic resolution and in the process elucidates its possible substrate-recognition mechanism.

## INTRODUCTION

Cytochrome P450s are key hemoprotein monooxygenase family which catalyzes a diverse range of biochemical processes, including metabolism of almost all drugs, lipid and steroid biosynthesis and decomposition of pollutants(1). They also play a pivotal role in the detoxification of xenobiotics, cellular metabolism, homeostasis and therefore, are of major clinical significance (2). These superfamily are especially notable for catalysing the hydroxylation process of unactivated hydrophobic molecules(3), making them attractive models for investigation of determinants of biological molecular recognition process(4–6). Over the years, cytochrome P450 systems from a variety of bacteria has found application in the development of an engineered catalyst for regio and stereoselective conversions of many compounds into useful products via the directed evolution approach.(7–9) Among various Cytochrome P450 family, CYP101A1, from the *Pseudomonas Putida* has served as the archetypal model for thermodynamic and kinetic investigation of substrate recognition in its heme centre(10,11). Popularly known as P450cam, this specific subfamily of cytochrome P450 catalyzes the hydroxylation process of D-camphor to 5-exohydroxycamphor(12,13). P450cam is known for its high substrate specificity in comparison to the specificities of many members of the P450 superfamily and this has been attributed to its relatively small, rigid active site that showed few binding-induced changes in early crystal structures(13). However, due to large attention from the community over last several decades, P450cam has been a regular subject of biochemical and crystallographic investigations, which has given rise to multiple reports of crystallographic structures in both substrate-free(14,15) and substrate-bound form of P450cam,(12, 13, 16). A visual inspection of these crystallographic conformations indicates that some of them deviates considerably from early perception of rigid active site of P450cam. As would be discussed in the forthcoming paragraphs, the prevalent heterogeneity in crystallographic structure of substrate-free P450cam has clouded our perception of underlying mechanism of substrate-recognition process in this protein.

The substrate recognition channel of P450cam is formed by F and G helices, the intervening F-G loop, and B’ helix, which fold around the heme and helix I to enclose the substrate and position it for binding of substrate to the heme (see figure 1).(12,17) The earliest crystallographic structure of P450cam in substrate-bound pose, resolved by Poulos and coworkers, revealed that the substrate is hidden in a deeply buried cavity near the heme active site, with no open channel accessible from protein exterior. (12) Discovery of crystallographic structure of a substrate-free conformation of P450cam (14) (pdb id: 1PHC) also indicated a substrate binding site that is not readily accessible from the outside (see figure 1A). Additionally, the conformations of P450cam describing the intermediate steps involving the catalytic pathways of this metalloenzyme have also been observed in occluded conformation.(18) The prevalence of these closed conformations in various states of activity of P450cam, including the precedences of a closed substrate-free conformations lead to a perception that during catalytic cycle, P450cam perhaps opens up to allow the camphor entry, closes the active site to perform the catalytic activity and then reopens for product release.(19)

**Figure 1:**
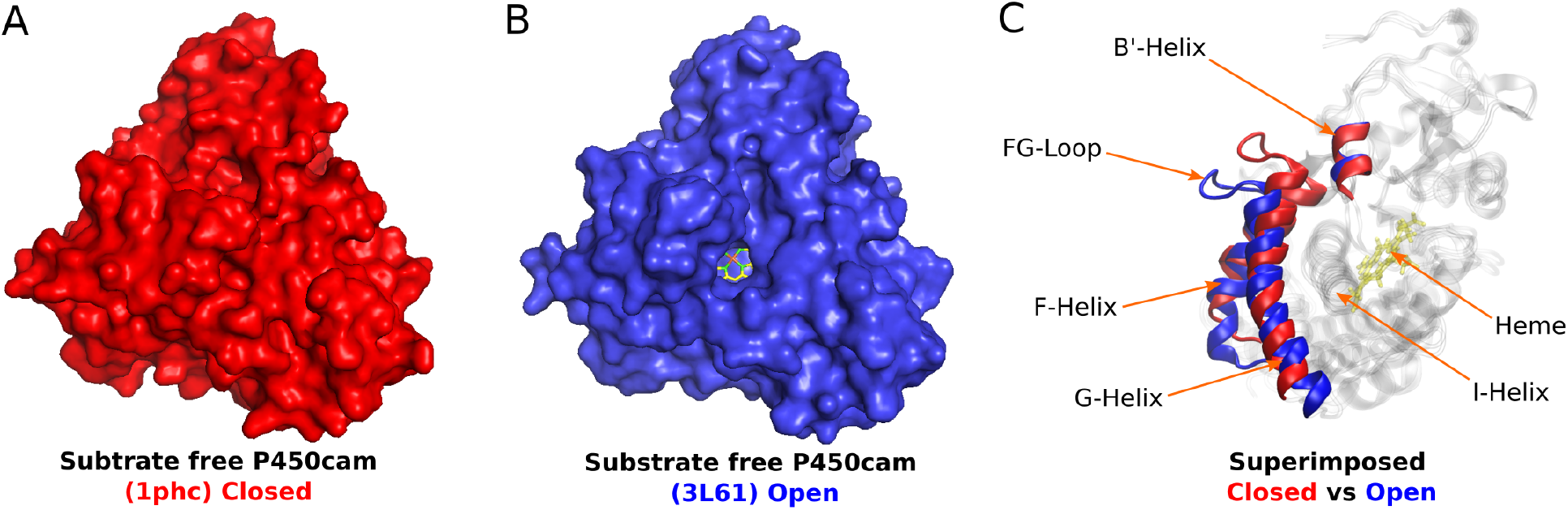
Systems under investigation: (A) Substrate free cytochrome P450cam (PDB:1PHC) with closed pocket (buried cavity). (B) Substrate free cytochrome P450cam (PDB:3L61) with open pocket (solvent accessible cavity). The heme is situated at the catalytic center and is visible in yellow-green color. (C) Comparison of Closed vs Open structures.

However, the precedences of occluded substrate-free crystallographic pose notwithstanding, report of first structure of an open conformation of P450cam in the presence of large tethered substrate analogs(20) and more significantly a more recent discovery of a crystallographic structure of ‘open’ conformation of substrate-free P450cam (pdb id: 3L61)(15) (see figure 1B) have complicated an otherwise simple picture of the conformational landscape of P450cam. Overlay of ‘open’ (pdb id: 3L61)(15) and ‘closed’ (pdb id: 1PHC)(14) crystallographic pose of P450cam shows that these two structures mutually differ only in the position of F/G helices and intervening F/G loop near the substrate binding channel (see figure 1C): The inward movements of F/G helices and F/G loop towards the heme active site are found to occlude the substrate-binding site in the closed conformation, while the outward movements of F/G helices and F/G loop away from the heme active site are found to open the substrate-binding site in the open conformation. Subsequent investigations of P450cam in solution by double electron-electron resonance (DEER) spectroscopy suggested that in the physiologically relevant solution state, the substrate-free P450cam exists in the open conformation (21) and it is the binding of substrate which leads to a closed conformation. Recent nuclear magnetic resonance (NMR) studies(22,23) showed that P450cam populates an ensemble of conformations based on the segmental movement of F and G helices. The presence of a mobile active site and speculation of conformational selection based substrate binding mechanism have been supported by Infrared (IR) spectroscopic investigations(24). Conformational fluctuation is known to play key role in substrate recognition across multiple P450s, but only recently have specific details about the conformational space sampled by a given enzyme are emerging(15, 25–27). Together, these findings have introduced the notion of complex ‘conformational heterogeneity’ in the substrate-free P450cam and its role in modulating substrate recognition by P450cam is slowly being appreciated. Additionally, the extent of structural changes traversed by a given enzyme and the kinetics between them is not well understood for any p450 enzyme(16). In this context, a quantitative analysis of conformational heterogeneity substrate-free P450cam at an atomistic resolution is in order and the current work performs this important task.

The current article quantifies the inherent conformational heterogeneity in P450cam by computer simulation of inter-state transition between ‘open’ and ‘closed’ conformation of substrate-free P450cam. The substrate free ‘closed’ structure of P450cam (PDB id: 1PHC)(14) and the substrate free ‘open’ structure of P450cam (PDB id: 3L61)(15) form the basis of our work (see figure 1). We explore the possibility of transition from one state to the other state, by spawning unbiased Molecular Dynamics (MD) simulation trajectories from the both end. We use an aggregated 84 *μs* of MD trajectories to reconstruct the underlying conformational landscape of P450cam and decipher both free energetic and kinetic aspects of conformational transition within the framework of Markov state model (MSM)(28–30).

In particular, our results show that for a substrate-free P450cam the MSM identifies the ‘closed’, ‘open’ and ‘intermediate’ clusters or ensembles of conformational states of P450cam in which closed state was obtained as a dominant conformation. Additionally, the exhaustive nature of the computer simulations also detects an interesting feature of uncoiling of the B’ prime helix of P450cam and identifies other two clusters or ensembles of conformations, which are found to be coupled to the open conformation(15,31). Although the mapping of all these conformational states on to the free energy landscape suggests these are separated by small or accessible energy barriers, the kinetic net flux information along with their stationary population statictics suggests following: while P450cam can in principle visit both open and closed conformation, the balance of the conformational equilibrium is tilted towards ‘closed’ conformation and a potential substrate of P450cam would mostly encounter a closed conformation, hence would follow an induced-fit based recognition process, as reported by Ahalawat and coworker(32). Interestingly, the kinetic flux network of transition paths also predicts that an intermediate ensemble facilitates the transition between open and closed conformation. These results reconcile multiple prior experiments(15, 16, 21–24) and simulations (32–34). Taken together, the current work renders a quantitative account of conformational heterogeneity in P450cam and provides key insights on its implication in possible mechanism of substrate recognition in P450cam.

## MATERIALS AND METHODS

### Unbiased atomistic simulations of substrate free P450cam

The unbiased atomistic simulations were performed using the substrate-free crystallographic poses of the proteins. Simulations were spawned from *both* the substrate free ‘closed’ (PDB id: 1PHC)(14) and ‘open’ structure of P450cam (PDB id: 3L61)(15). Both of these structures have 99.9% sequence similarity except at 334th position where 1PHC has Cystine(CYS) residue and the 3L61 has an alanine (ALA) residue, leading to only one atom difference. We therefore computationally mutated cystine to alanine in pdb id 1PHC for comparison of these two crystal structures at equal footing. Further, we modeled the missing coordinates of B’ helix in the 3L61 starting structure using the homology modelling program Modeller(35) by using 1PHC as a template.

Charmm27 force fields (36) were employed to model the protein and the heme segment in their all-atom representations. In each cases, a 8.9nm × 8.9nm × 8.9nm cubic box was solvated with TIP3P(37) water, with the protein centred at the box, with periodic boundary condition implanted in all three directions. Sufficient numbers of sodium and chloride ions were added to keep the sodium chloride concentration at 150 mM and render the system charge neutral. The total number of particles in each system was around 71000.

All simulations were performed in NPT ensemble using leap-frog integrator with 2 fs time step. The Verlet cutoff scheme(38) was employed throughout the simulation with The Lenard Jones interaction extending to 1.0 nm with dispersion corrections. The electrostatic interactions were implemented with a short-range electrostatic cutoff at 1 nm, while the long range interactions were treated by Particle Mesh Ewald summation(39) with cubic interpolation and in Fourier-grid space of 0.16 nm. The neighbour lists were updated every 10 steps. All bond lengths involving hydrogen atoms of the proteins and the ligands were constrained using the LINCS algorithm and water molecules were kept rigid using the SETTLE(40) approach. The average temperature is maintained at 300 K by coupling protein and water separately to a Velocity rescale thermostat(41) with a relaxation time constant of 0.1 ps. The average isotropic pressure of 1 bar was maintained using Parrinello-Rahman barostat.(42) with a relaxation time constant of 2 ps and isothermal compressibility of 4.5 × 10^5^ bar^-^1. The simulation trajectories were saved every 10 ps for the analysis. All simulations were performed using GROMACS 2018 (43).

Figure 2 provides the schematic of the overall methodology adopted in this work. We first quantified the ‘open’ and ‘closed’ conformations based on the collective root-mean squared deviation (RMSD) of F/G-helices plus intervening FG-loop region and B’-helix relative to ‘closed’ crystallographic pose. Towards this end, we aligned the MD trajectories with the ‘closed’ crystallographic pose along the backbone residues of the I-helix (resid 234 to 267), “B5”-beta sheet (resid 389 to 401), B’-helix (resid 89 to 96) (see figure 3A) via rotation and translation and subsequently calculated the collective RMSD of F/G-helices plus intervening FG-loop (resid 173 to 214) and B’-helix region (resid 89 to 96) (see figure 3B). As discussed in the next paragraph, extensive MD simulations were undertaken to quantify the underlying conformational landscape of P450cam and to develop a Markov State Model (MSM) for describing the underlying dynamical process.

**Figure 2:**
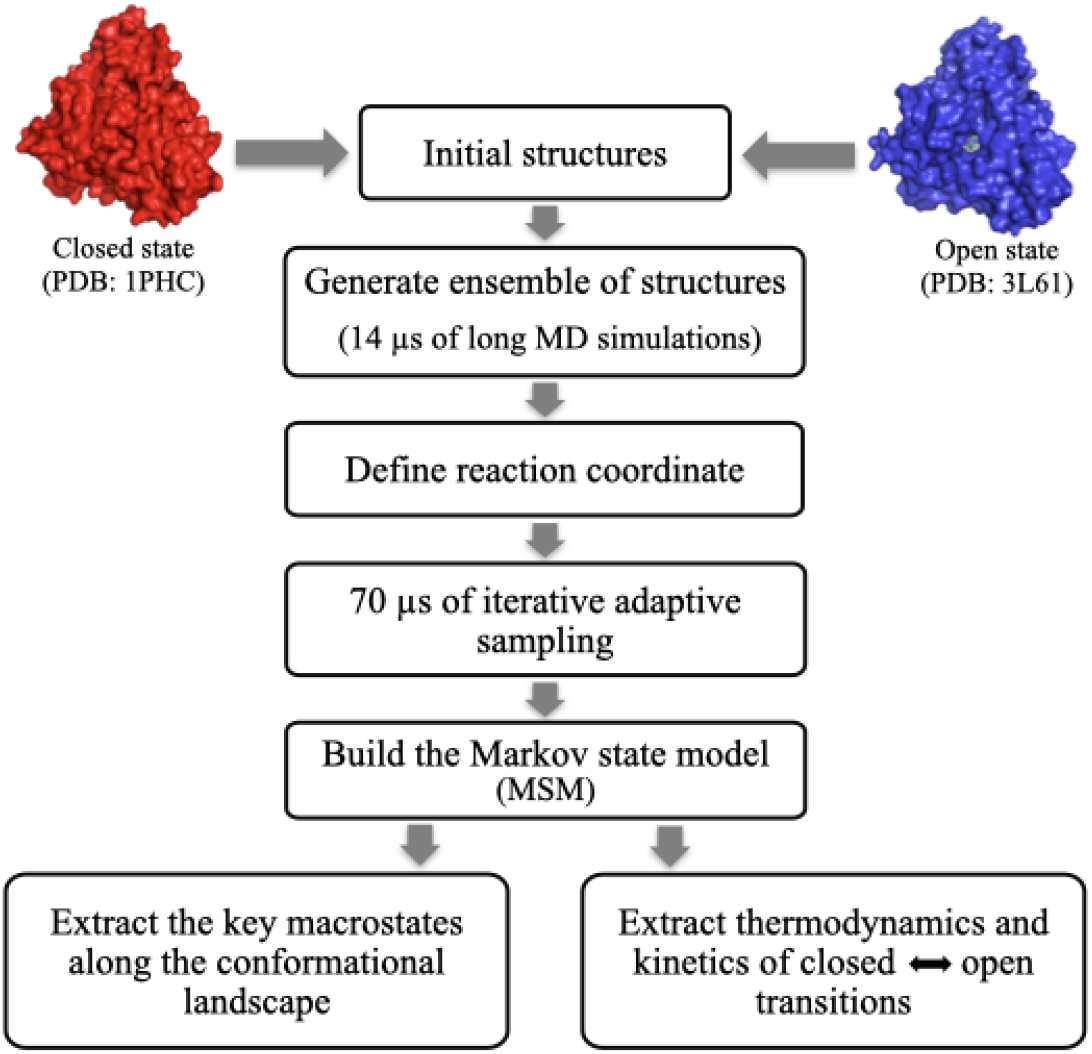
The schematics of the overall methodology adopted to capture conformational heterogeneity in cytochrome P450cam system.

In the current work, we have adaptively sampled the conformational phase-space of the substrate-free P450cam. Figure S1 schematically illustrates the adaptive sampling approach employed in this work. Towards this end, we first initiated a series of long unbiased MD simulations of substrate-free P450cam from both ‘closed’ crystallographic pose (pdb id: 1PHC) and ‘open’ crystallographic pose (pdb id: 3L61) by generating random initial velocities according to Maxwell-Boltzmann distribution. From each initial configurations, 14 trajectories of each 500 ns length were initiated leading to 7 μs (i.e 14 × 500 ns) of MD simulations. Hence a total of 14 μs long unbiased MD simulations from both the initial configurations were performed. On each of the long simulations data set from ‘closed’ and ‘open’ conformation we then performed the k-means clustering by projecting the data along the chosen CVs (CV choices would be justified in the upcoming paragraph) to generate 200 clustercenters, each of which served as the seed of the additional simulations conducted for adaptively sampling of the conformational phase space. Subsequently, independent relatively shorter MD simulations of 100 ns length, with random velocity seeds, were spawned from each cluster centre. Some of the simulations corresponding to a region where RMSD1 ≥ 0.2,RMSD2 ≥ 0.2 (the two CVs chosen in the present work and discussed later) were extended till 250 ns to improve the sampling in this region. The trajectories so obtained in this iteration were re-aggregated, further projected in CV space and re-clustered via regular space clustering. The data-poor regimes in this phase-space, identified from this iteration, were subjected to a new series of short MD simulations. The adaptive sampling approach employed this way exhaustively covered the conformational landscape. As shown by a fully connected reversible kinetic network in SI (discussed later in MSM part), the finite non-zero values of rate of bi-directional interconversion among all macrostates suggest that the adaptive sampling employed here has covered possible transitions among key states in the conformational landscape.

### Markov state model analysis

We undertook the task of building a Markov state model (MSM)(29, 30) for gaining the detailed understanding of conformational heterogeneity in P450cam using 84 μs data of unbiased all-atoms simulations. We employed PyEMMA (44)(http://pyemma.org) for the purpose of constructing the MSM.

Prior to construction of MSM, we first assessed and validated the choice of optimal CVs for describing the open/close process of P450cam. A good CV should be able to demarcate the key open and close states in a precise and quantitative fashion. The visual inspection and preliminary analysis of the trajectories indicated that, apart from the opening and closing events of F/G-helices and intervening F/G-loop of P450cam, uncoiling of B’-prime helix is also taking place as an additional dynamical event. We therefore defined two separate collective variables (CVs) for this purpose: RMSD of F/G helices named as *RMSD*_1_ (using residue id 173 to 185 and 192 to 214 only) and RMSD of B’-helix named as *RMSD*_2_ ( using residue id 89 to 96) (refer figure 3B). We omitted the intervening F/G-loop region in the *RMSD*_1_ calculation because of its higher flexibility to decrease the noise in the calculation. Before calculating these RMSDs we aligned the protein backbones of the I-helix, B’-helix and “B5”-beta sheet of all frames of the trajectory with the backbone of ‘closed’ crystallographic pose. Both these RMSDs were computed relative to ‘closed’ crystallographic pose. As shown in figure 4, these two RMSD-based CVs were able to demarcate a closed conformation from an open conformation.

**Figure 3:**
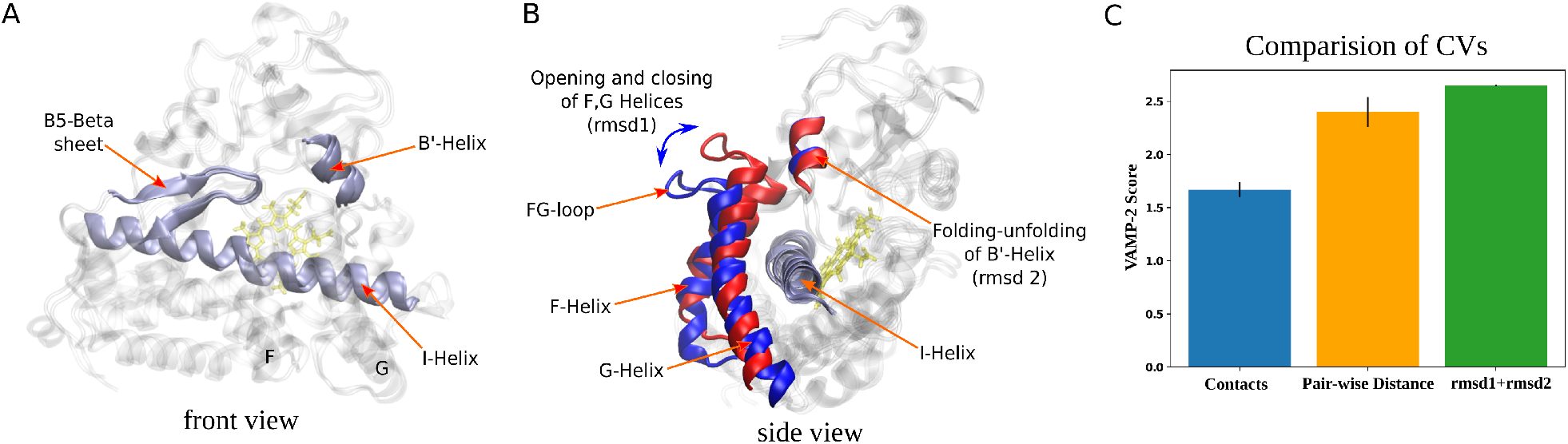
Definition of the reaction coordinate for capturing conformational heterogeneity in cytochrome P450 system: (A) The alignment region used for fitting in rmsd calculations. (B) The reaction coordinates chosen (*RMSD*_1_,*RMSD*_2_). In the figure A the protein is shown from the front side and FG-helices are kept transparent while in the figure B the protein is shown from the side view after rotating figure A by 90° along z-axis and the B5 sheet is kept transparent for clarity. (C) Comparison of multiple CVs such as number of native contacts, pair-wise distances and RMSDs based on the VAMP-2 score.

**Figure 4:**
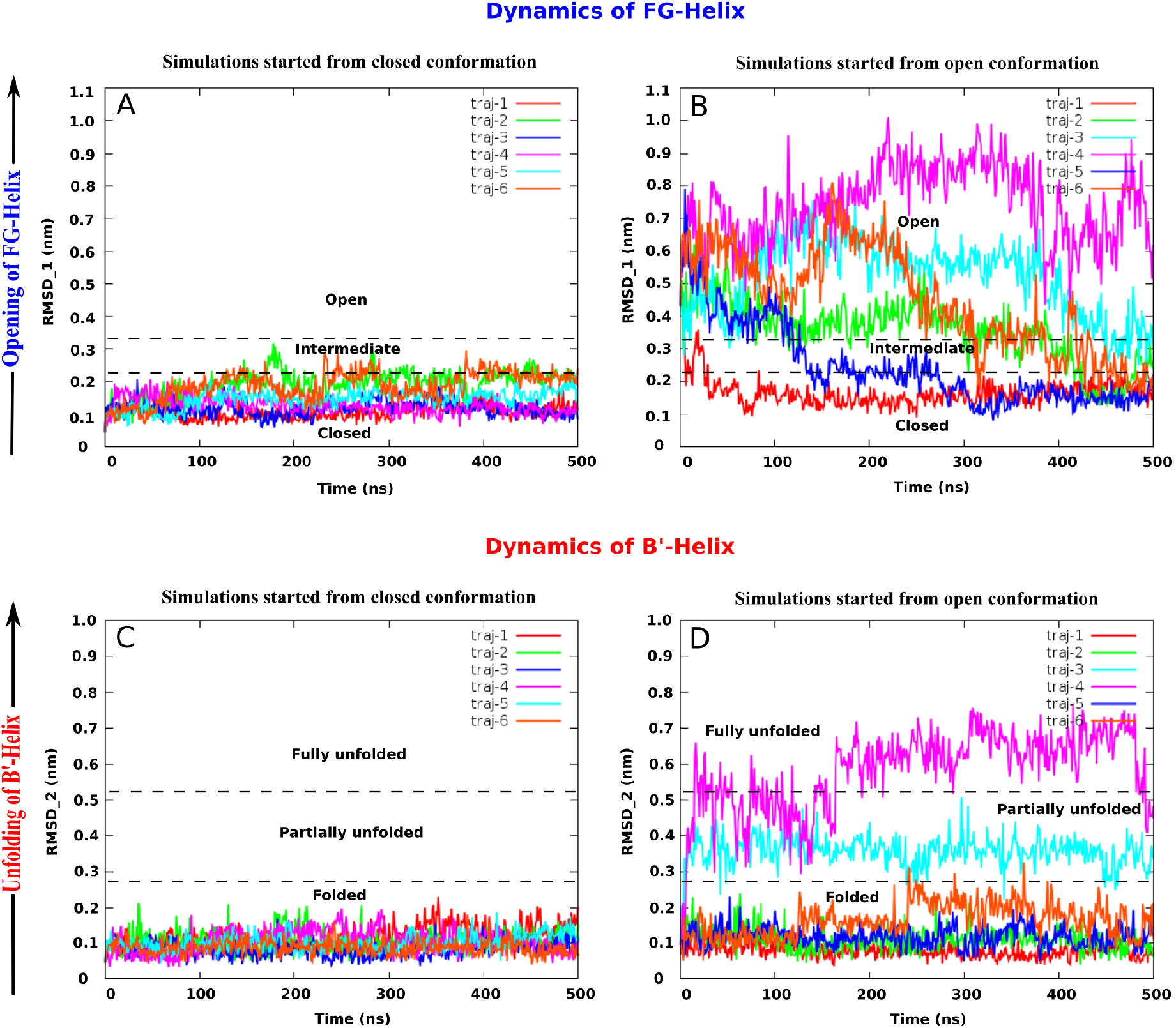
The conformational dynamics of FG-helix and B’-helix in cytochrome P450: (A) The time profile of *RMSD*1 for simulations starting from closed conformation and (B) for simulations starting from open conformation. (C) The time profile of *RMSD*_2_ for simulations starting from closed conformation and (D) for simulations starting from open conformation.

We compared the optimality of the chosen CVs with multiple potential candidate CV by comparing their respective VAMP-2 score.(45,46) As introduced by Noe and coworkers, the VAMP-2 score is based on a variational approach for Markov processes (VAMP) that allows one to find optimal features (CVs) and to cross-validate the hyper-parameters and to optimize a Markovian model of the dynamics from given time series data(45,46). Apart from RMSD-based CVs, we explored two other candidate CVs: pair-wise distance and number of native contacts. In case of pair-wise distance calculation we selected C-alpha atoms of I-helix, B’-helix, “B5”-beta sheet and F/G helices and intervening FG-loop regions and then calculated the distances for the pairs of selected CA atoms which are farther than 3 residues apart. To calculate the number of native contact calculation, we used similar selection of C-alpha atoms of I-helix, B’-helix, “B5”-beta sheet and F/G helices and intervening FG-loop regions. As shown in the figure 3C, we found that VAMP-2 score is relatively higher for (RMSD1,RMSD2) than the other two CVs.

Finally, we checked if time-structured independent component analysis (TICA)based projection of RMSD1 and RMSD2 would have provided any superior projection compared to the raw CVs. Accordingly, we performed TICA of RMSD1 and RMSD2 for various TICA lag times and compared the free-energy-surface (FES) projected along RMSD1 and RMSD2 to that projected along two TICA coordinates of RMSD1 and RMSD2. The FES obtained along the raw dimension and that along TIC dimensions are depicted in figure S2. We find that there is essentially no change in the FES when projected along TICA dimensions, apart from the fact that the FES gets rotated by a certain angle in (TIC1,TIC2) space. All other features are almost same in both cases. This suggests that in this particular case, FES along TICA is not any superior, when compared with FES along raw CVs. Accordingly, for better interpretability and more physical picture (which is otherwise lost in a projection or linear combination as in TICA), we use RMSD1 and RMSD2 as the optimal choice of CVs. Taken together via visual inspection of the crystallographic poses, comparison of VAMP-2 score and cross-checking the advantages of using TICA, we validated that RMSD1 and RMSD2 are optimal choices for demarcating the key states and for providing dynamically slow projections.

Accordingly, we used the (RMSD1,RMSD2) as clustering metric for discretising the simulation trajectories for subsequent MSM analysis. As shown in figure S3, VAMP-2 scores for a wide range of cluster numbers ranging from 100-2000 were very similar. We chose the highest number of cluster centers (2000) to get both kinetic and geometric resolution in the data. We employed the k-means clustering algorithm(47) to discretise the input data into 2000 clusters. Detailed balance is imposed on the MSM by symmetrizing the count probability matrix before matrix digonalization and is a standard step in MSM construction approach which is employed using PyEMMA. Then a 2000-microstate MSM was constructed at a lag time of 20 ns. The choice of lag time was based upon the onset of levelling of implied time scale, which ensures the Markovianity of the model (see figure 5B). The implied time scale plot suggested the presence of total 5 metastable states at 20 ns lag time. The 2000-microstate MSM was coarse-grained into a 5 macrostate model using Perron Clusters Cluster Analysis method (PCCA) (48). The MSM is built with 95% of confidence interval and all the values calculated from the model are with the same level of accuracy. We also performed the bootstrapping of the MSM analysis to estimate the standard error in the stationary populations of all five macrostates.

**Figure 5:**
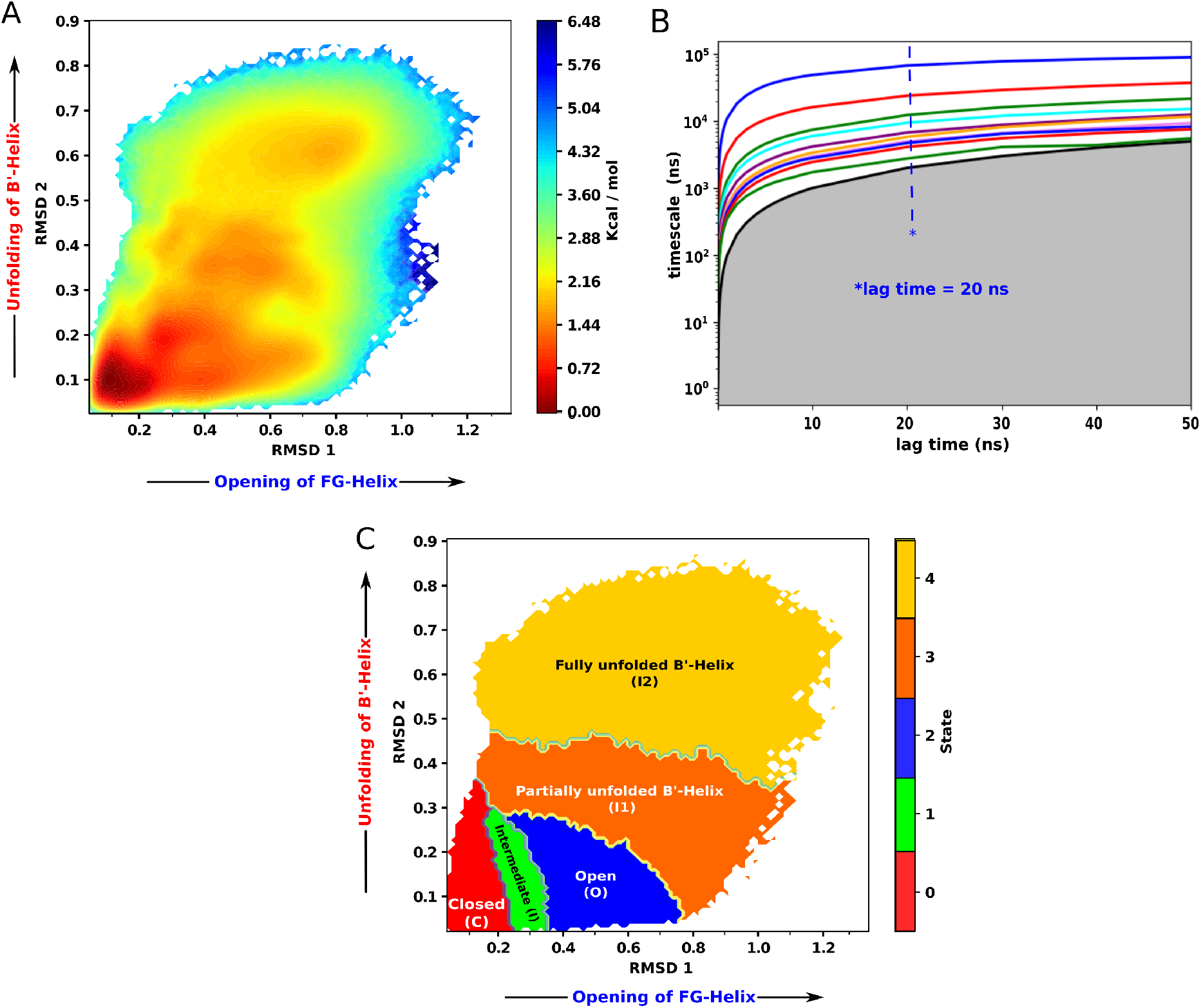
The thermodynamics of the conformational heterogeneity in P450cam: (A) MSM reweighed free energy landscape. It shows a presence of five distinct energy basins along *RMSD*1, *RMSD*2. (B) The implied time scale (ITS) plot of the constructed Markov state model. (C) The state decomposition obtained from MSM calculation. Each macrostate shown in different colour and labelled with the description of conformations it represents.

The mean first passage time (MFPT) for transition between two states was computed from the MSM. The MFPT defines an average timescale for a stochastic event to first occur(49) or the average time taken for a given conformation to fold for the first time (50). The MFPT, *F_ij_*; for the transition of state *i* to state *j* is given by the following expression(50)

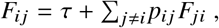

where (τ) is the lag time used, *p_ij_*; is the probability of transitioning from state i to state *j* within the lagtime of (τ) and the *F_ji_* is the MFPT for the reverse transition. The rate constants were calculated as

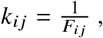

where *F_ij_*; is the MFPT for the transition of state *i* to the state *j*.

The gross flux, the average number of the transitions from state i to state *j* were estimated form the Transition Path Theory (TPT)(44, 51–53) and are computed as follows

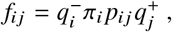

where 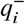 and 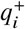 are the backward and forward commitor probabilities, *π_i_* is the equilibrium population of the state and *π_i_p_ij_* is the equilibrium flux. The net fluxes are computed as

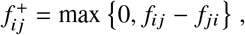

These net fluxes define a directed path from initial state A to final state B and hence define the mechanism of the dynamics between A and B.

## RESULTS

Precedences of crystallographic poses of both closed and open structures of P450cam in its substrate-free form raise a question if these two crystallographic poses are dynamically visited by P450cam in the array of conformational ensemble. A quantitative insight into this question would address mechanistic aspect of recognition process of camphor to P450cam: For example, a prevalence of closed conformation in the conformational ensemble of P450cam would lead to an induced-fit based mechanism for substrate recognition by P450cam. On the other hand, prevalence of open conformation or equal probability of both open and closed conformation in the conformational landscape would imply that the substrate might adopt a conformational selection based mechanism for recognising P450cam. To resolve this question, we first simulate the conformational dynamics of substrate-free P450cam by initiating a large set of MD trajectories from both open (pdb id: 3L61) and closed (pdb id: 1PHC) crystallographic poses. Figure 2 illustrates the schemes of overall methodology adopted in the present work (see method for details). An overlay of the crystallographic pose of open and closed conformation (see figure 1C) suggests that these two structures mutually differ by an RMSD of 0.45 nm due to difference in positioning of F/G helices region. Accordingly, we monitor the root-mean squared deviation (RMSD) of F/G helices of all trajectories from that of closed conformation (pdb id: 1PHC) (hereby referred as *“RMSD*_1_”, see methods section and figure 3 for illustrative details of the metric employed) as a measure of the conformational dynamics of P450cam.

We first compare the representative time profiles of RMSD of F/G helices as computed from equilibrium simulated trajectories, initiated from closed and open crystallographic pose respectively (figure 4A and 4B). We find that for trajectories starting from the open crystallographic pose (refer figure 4B), *RMSD*_1_ decreases from 0.45 nm to 0.20 nm (or below that) at a very short time scale, implying that the ‘open’ conformation of P450cam often makes transition to closed conformation within the simulation time scale. In some of the trajectories initiated with open conformation, we also find that *RMSD*_1_ increases from 0.45 nm to 0.9 nm within a very short time scale, implying that the ‘open’ conformation of P450cam potentially can make transition to even more open conformation also within the simulation time scale. On the other hand, none of the MD trajectories starting from the closed crystallographic pose (refer figure 4A) made transition to open conformation within the simulation period. Rather, in trajectories initiated from closed crystallographic pose, *RMSD*_1_ fluctuates between 0.04 and 0.20 nm, within the simulation time scale, at times making occasional visit to an intermediate state(*RMSD*_1_ ≈ 0.22 nm), implying that trajectories initiated from closed crystallographic pose mostly remained confined around closed conformation within the time-scale of the simulation. Inspection of simulation trajectories initiated from open conformation also reveals unfolding of B’ helix in some cases. As depicted in figure 4D, RMSD of B’ helix (referred as *RMSD*_2_, see figure 3B) deviates significantly in trajectories initiated from open conformation. However, the *RMSD*_2_ does not change substantially in trajectories initiated from closed conformation and the B’ helix remains helical. (see figure 4C). These observations are also evident from the free energy surfaces (FES) shown for long simulation data obtained from close and open state.(see figure S1 A,B)

Taking cue from the trends of conformational transitions in these simulation trajectories, we reconstructed the underlying conformational free energy landscape of P450cam to get detailed insights of complex conformational dynamics. Towards this end, we combined long simulation trajectories initiated from both ends (closed and open crystallographic pose, see figure S1 E) and adaptively sampled conformational space by spawning numerous short simulations from various conformations clustered from the long trajectories (see figure 2 and figure S1 for illustration of adaptive sampling schemes). The overall simulation efforts resulted in a 84 *μs* conformational ensemble of substrate-free P450cam (see methods). These aggregated trajectories were then discretised and a Markov state Model (MSM) was built and a free energy landscape along *RMSD*_1_ and *RMSD*_2_ is derived. This free energy landscape is then finally reweighed according to MSM-derived relative stationary weights for each microstate(54).

Figure 5A depicts the two-dimensional map of the free energy landscape along *RMSD*_1_ and *RMSD*_2_ and provides a quantitative thermodynamic account of underlying conformational heterogeneity in P450cam. We find that the location of global minimum of the free energy landscape corresponds to both *RMSD*_1_ and *RMSD*_2_ close to 0.1 nm, which is very close to the RMSD of the ‘closed’ crystallographic pose (pdb id: 1PHC). Apart from this, an important basin at a relatively higher free energy value, centred around *RMSD*_1_ = 0.45 nm and *RMSD*_2_ = 0.10 nm, corresponds to the ‘open’ crystallographic pose of pdb id 3L61. These two minima corresponding to closed and open conformation were estimated to be separated by a small free energy barrier is 1.0 kcal/mol. Additionally, the free energy surface also showcases three other local minima at other values of *RMSD*_1_ and *RMSD*_2_. One of the minima lies in between open and closed state (*RMSD*_1_=0.3 nm and *RMSD*_2_=0.20 nm), with the small free energy barrier from both the side of about 1-1.2 kcal/mol which we would eventually identify as an intermediate macrostate. The remaining two minima belongs to relatively larger value of *RMSD*_1_ and *RMSD*_2_: one with partial unfolded B’-Helix corresponding to *RMSD*_1_ and *RMSD*_2_ values of 0.55 nm and 0.38 nm respectively and other with fully unfolded B’ helix corresponding to *RMSD*_1_ and *RMSD*_2_ values 0.78 and 0.62 nm respectively. Again, these states are separated from other states with the accessible barrier of about 2.0 and 2.2 kcal/mol respectively. Together, the presence of multiple basins at different free energy values with small or accessible free energy barrier reflects a conformationally heterogenous landscape of substrate-free P450cam. Overall, our calculations were able to obtain the ‘closed’, ‘open’ and ‘intermediate’ ensembles or clusters which are consistent with the prior experimental investigations.(16,22)

To quantify the state-space decomposition of the conformational landscape in more comprehensive manner, the microstates constituting the underlying MSM are coarse grained into finite number of macrostates. The implied time scale plot as a function of lag time (figure 5B) suggested that a 5-state decomposition at 20 ns lagtime would be optimal for aptly describing the conformational landscape of P450cam. We note that the separation of time scales are smaller beyond 3 macrostates. However, our observation suggests that in an otherwise 3-state decomposition, the open state is found to be a mixture of conformations with folded, partially folded and fully unfolded B’ helix. Accordingly, for a more fine-grained discretisation of the open conformations, we construct a 5-state MSM in a bid to resolve the complex B’-prime unfolding dynamics. The smaller gap in the implied time scale plot beyond 3 states also reflect that these states are kinetically not very separated. As we would find later, the 5-state decomposition would reveal those states to include conformations with partially folded and fully folded B’ helix. We find that each of the five macrostates (namely C, O, I, I1 and I2) corresponds to a distinct basin of the free energy landscape along *RMSD*_1_ and *RMSD*_2_ (figure 5C). Table 1 enumerates relative stationary populations of these macrostates. We find that the closed conformation (C) is the most populated (54.67 %) macrostate and is spatially well-separated from the open conformation (O) (19.59 %) via an intermediate conformation (I) (11.60 %). Interestingly, apart from identifying the ‘closed (C)’, ‘open(O)’ and intermediate (I) states, we also discover two additional macrostates (I1 and I2) of relatively lower populations at larger values of *RMSD*_1_ and *RMSD*_2_.

**Table 1:** The stationary populations for MSM macrostates.

| MSM<br>macrostate | Populations<br>in % |
| --- | --- |
| State C | 54.67 ± 0.05 |
| State I | 11.60 ± 0.06 |
| State O | 19.59 ± 0.04 |
| State I1 | 8.48 ± 0.01 |
| State I2 | 5.64 ± 0.02 |

In figure 6, we depict distinct five representative snapshots corresponding to most probable states from each macrostates. These conformations are also annotated on the free energy landscape in figure S4. Each of the snapshot consists of ten conformations randomly pruned from ensemble of each macrostates and overlaid with the closed crystallographic pose (1PHC, red) for comparison. Visual inspection of these conformations (refer figure S4) shows that there is gradual opening of the F/G helices from close conformation (C) to the intermediate conformation (I) and to the open conformation (O). In all these three states the helicity of B’-helix is preserved. The overlay of conformations of macrostate C, macrostate O and macrostate I in figure 7 (left) clearly shows that the state-space decomposition can distinguish the conformations based on the extent of outward movement of F/G helices (away from heme group) from the closed crystallographic pose: the F/G helices move the most, away from heme group, in macrostate O, followed by that in macrostate I and least in macrostate C. Visual inspection of the conformations corresponding to I1 and I2 indicates that these two macrostates appear due to occasional uncoiling or unfolding of B’ prime helix as F/G helix opens. The overlay of conformations of macrostate I1 and macrostate I2 with the macrostate C (refer figure S4) indicates partial to complete uncoiling of B’ helix, with outward movement of F/G helices in these conformations resembling that of macrostate O.

**Figure 6:**
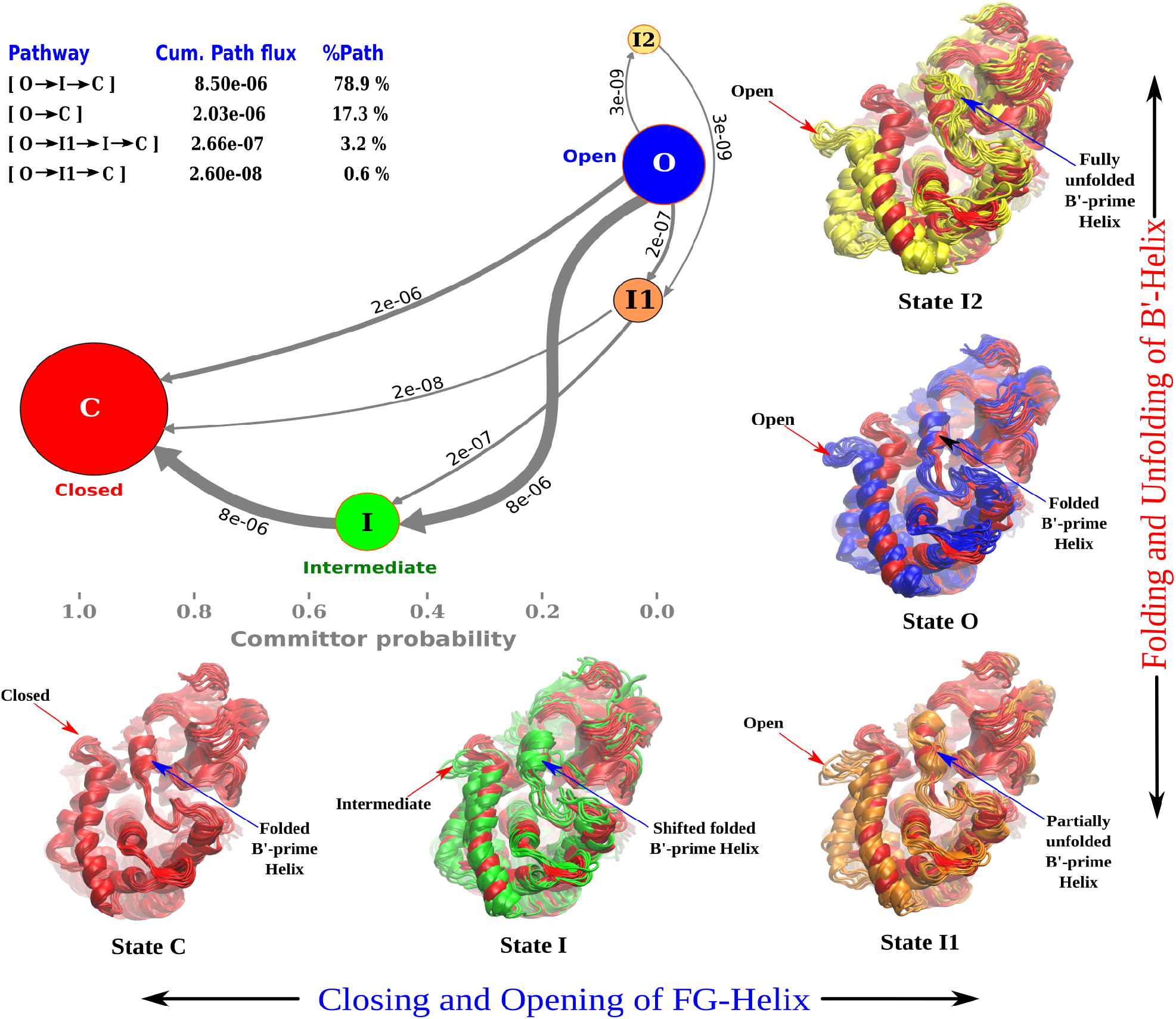
The kinetic mechanism of conformational heterogeneity in cytochrome P450cam: The net flux network is obtained from TPT analysis. The network connects five macrostates and are represented as a circular discs. The sizes of the discs are proportional to their stationary populations and are placed according to their respective committor value. The committor scale is drawn at the bottom of the figure. The committor value of 1 corresponds to the closed state(C) and 0 corresponds to the open state(O), while committor value of 0.5 corresponds to the intermediate (I) state. The states are connected with the arrows and thickness of the arrows is proportional to the amount of net flux of the transition within the macrostates. The flux values of corresponding transitions are written over the arrow. The resulting pathways are shown at the top left of the figure along with their cumulative path fluxes and percentage path contribution. The figure also shows the distinct five representative snapshots corresponding to most probable states from each macrostates. The snapshots are shown in comparison with the first state, the closed state C (red colored) to indicate the relative differences in the conformations. In each states 10 snapshots are shown for simplicity.

To unravel the underlying kinetic mechanism of conformational heterogeneity in cytochrome P450 we constructed the kinetic network of transition among the macrostates, using Transition Path Theory (TPT). We considered closed state(C) and open state(O) as two extreme ends of the conformation that P450cam chooses to explore and extracted the net fluxes of the transitions among the macrostates. Figure S4 depicts that the conformational transitions are bi-directional between any pairs of macrostates, indicative of exhaustive sampling of conformational phase space. However, the extent of flux is not same in both the directions. Figure 6 depicts the kinetic mechanism of conformational transitions in substrate-free P450cam based on the direction of the net flux and the amount (arrow thickness) of net flux among the macrostates. These macrostates are represented as a circular discs with their sizes proportional to their stationary populations and are placed according to their committor probability value. The kinetic network (figure 6) dictates that while there are finite probabilities of transitions between open and closed state on both directions, the direction of the net flux arrows dictates that the overall transitions will be from O → C, implying that the net equilibrium of (*C* ⇌ *O*) transitions would be tilted in favour of closed (C) conformation. The MFPT values for state-to-state transitions depicted in figure S4 also support that rate of *C* → *O* transition is slower than the reverse process *O* → *C* transition. The arrow thickness in figure 6 indicates the amount of net flux among the macrostates. Figure 6 also depicts that there are *multiple* pathways for transition between the closed (C) and open states (O). Especially, comparison of net flux values among different pathways dictates that the *direct* transition *O → C* is less likely, with only a 17 % contribution to the total path. On the other hand, the net flux values predict that the transition from open (O) state to closed (C) state would be predominantly mediated by intermediate macrostate (I). In particular, O → I → C would emerge as the most dominant pathway with a 78.9 % of contribution in the total path. The kinetic network also suggests additional pathways of transition mediated by I1 ((O → I1 → I → C) and (O → I1 → C)), but their contributions in the overall pathways are very small. Taken together, higher stationary populations of closed macrostate (table 1), coupled with faster rate (figure S4, table 2) and net flux (figure 6) of open (O) to closed (C) transition (with or without the intermediate(I)) unambiguously indicate that the dynamic equilibrium of conformational transition is tilted in favour of closed conformation of P450cam. This is also schematically shown in figure 7 (right).

**Figure 7:**
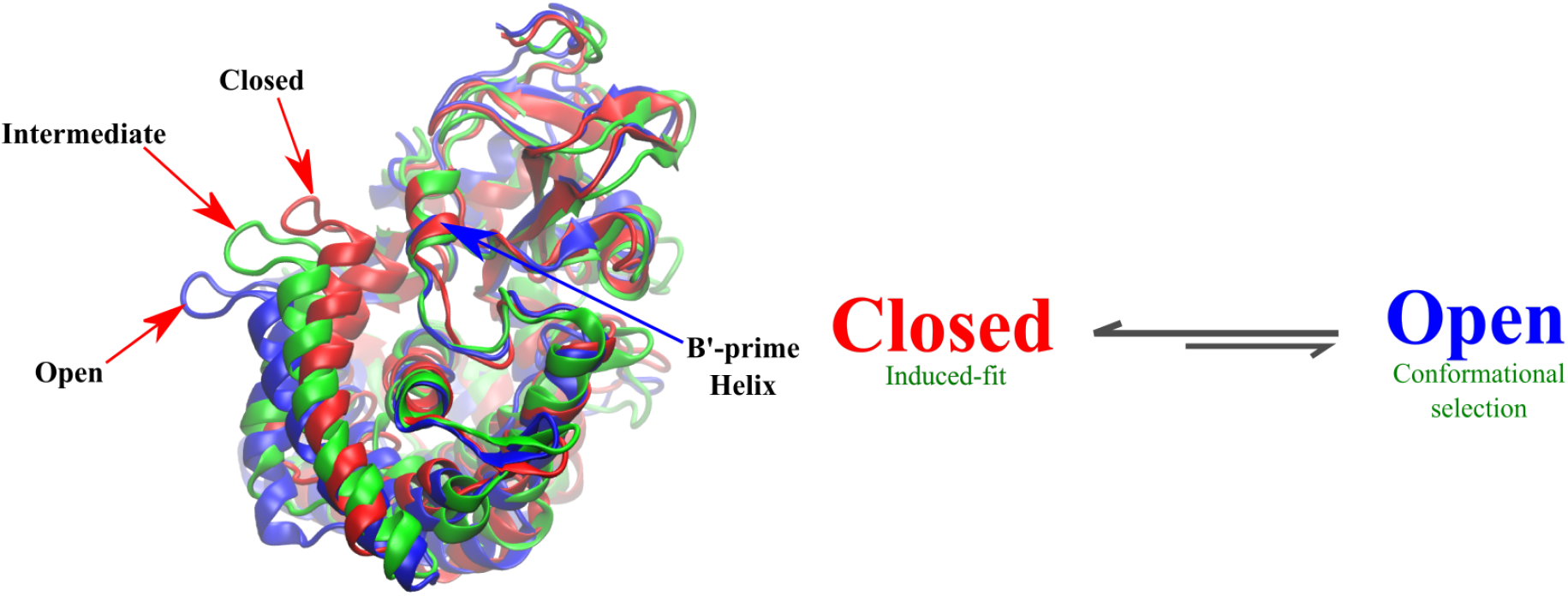
Left: Comparison of major MSM macrostates, macrostate C (closed), macrostate I (intermediate) and macrostate O (open). In all these three states the helicity of B’-helix is preserved. Right: Proposed substrate recognition mechanism

**Table 2:** The kinetic rate of mutual transitions between state C and state O.

| State<br>transitions | Kinetic constants<br>in $s^{-1}$ |
| --- | --- |
| State C → State O | $k^{co} = 4.13 \times 10^{-4}$ |
| State O → State C | $k^{oc} = 20.81 \times 10^{-4}$ |

Figure 6 further quantifies the respective committor probabilities of each of the five macrostates. We find the closed state (C) and the open state(O) are located at a committor probability of 1 and 0 respectively. Most interestingly, the intermediate (I) state corresponds to a committor probability of 0.5, suggesting that it is the saddle point in the transition path (O → I → C) with 50% (or equal) transition probability for both the direction. This is also consistent with the relative location of intermediate (I) macrostate in the middle of closed (C) and open (O) state in the free energy surface (see figure S4). The committor probabilities of intermediate states (I1) and (I2) are 0.02 and 0.01 relative to closed state, which also indicates that they are closely associated with the open state (O). This is consistent with the longer MFPT values for C → I1 transitions than O → I1 or O → I2 transitions (see figure S4) and suggests that there is very small probability of unfolding or uncoiling of B’-helix in closed states. Taken together, the observation of macrostates I1 and I2 in the conformational ensemble correlates very well with the partial to complete uncoiling of B’ helix with the outward movement of F/G helices.

In addition, figure S5 depicts the key interactions present across the MSM macro states from closed to open states. Figure S5A,B identifies key inter-residue interactions in the closed (C) state which are stabilising the interactions between F/G-helices residues with the I helix and BC-loop residues. These similar interactions are present in intermediate (I) state also but they gradually decrease from closed (C) to intermediate(I) state and to open (O) state. In open (O) state the interactions of F-helix with I helix are found to be broken, leading to increase in flexibility of overall F/G-helices (figure S5C). In I1 and I2 states (figure S5D) these interactions are completely vanished. More specifically, the loss of interactions of G-helix residues with BC-loop in I1 and I2 state causes increase in flexibility of BC-loop which leads to increase in fluctuation of B’-helix and makes it to unfold.

The observation of macrostates with partially or completely unfolded B’ helix in open conformation (i.e. I1 and I2 macrostates) is consistent with prior crystallographic investigations(15,16). In prior investigations by Lee et. al.,which involved crystallisation of the proteins in presence of camphor with low and high potassium ion concentration(15) or comparison of the crystallised structures of cytochrome P450cam bound to library of active substrates (16), the B’ helix has been found to provide key contacts with the substrate. On the contrary, in absence of substrate, cytochrome P450cam was found in open state where the B’ helix was in disordered (unfolded) state along with loss of its site specific potassium ion interaction due to loss of its interaction with the substrate(15,16). On a related note, it is also found that the B’ helix regains its folded state in the presence of substrate and finally converts to the closed state (PDB id: 3L63) (15,16). These experimental observations also justify why the closed substrate-free conformation (pdb id: 1PHC) is structurally identical with substrate-bound closed state, is more stable and is always associated with a folded state of the B’-helix.(15)

The functional implication of the current results can be assessed by exploring the most relevant question: How does the conformational landscape of P450cam, as portrayed in the current work, impact the substrate recognition mechanism by P450cam? While the current work predicts a conformationally heterogeneous landscape of substrate-free P450cam, a considerably larger population of ‘closed’ conformation than ‘open’ conformation, higher net fluxes from *open* → *closed* via intermediate coupled with the faster rate of *open* → *closed* transition, would tilt the conformational equilibrium towards the ‘closed’ conformation. As a result, we infer that a potential substrate like camphor would mostly visit the more populated ‘closed’ conformation of P450cam, which would require an induced-fit based mechanism for substrate-recognition (see figure 7, right for a schematic). This inference of ‘induced-fit’ based recognition mechanism is in line with Ahalawat et al’s recent investigation(32) of substrate-recognition process by P450cam. In Ahalawat et al’s multi-microsecond long simulation starting with ‘closed’ conformation of P450cam, camphor first diffused from bulk to the F/G helices of the protein and then spent a significant time to induce an opening, before landing on the designated heme active site. Similar to current work, in their investigation also, the F/G loop of the ‘closed’ conformation did not open up completely during multi-microsecond long trajectories. This induced fit mechanism of camphor binding to closed cytochrome P450 is also observed in the recent coarse grained simulations studied by Dandekar et. al. (55) where they observed binding of coarse grained camphor to a coarse grained CYP450 protein by inducing subtle changes along binding pathways in the protein without involving major conformational change. Recent kinetic evidence of an induced fit based mechanism(56) for substrate recognition in P450cam also provides strong support in this regard. However, the current work’s observation of a dynamical equilibrium between closed and open conformation (with the equilibrium being tilted towards closed conformation) also does not completely rule out the possibility of ‘conformational selection’ based substrate-binding mechanism, in which the substrate can potentially visit an open conformation and then closes the pocket after the binding is complete. (16,21)

## DISCUSSION

### Summary and implication of current work

Conformational heterogeneity plays important role in defining the structural and functional dynamics of the enzymes. In Cytochrome P450s, the identification and characterisation of the potential channels connecting its active site to the protein exterior is very important and the conformational heterogeneity has a critical role in their functional mechanism. It has also become apparent that the binding mechanism of substrate is linked to the enzyme’s ability to access its conformational ensemble in the presence and absence of substrate. The static three-dimensional crystallographic structures of enzymes solved in different conditions and/or environments are crucial to provide the conformational substates of enzymes but are not sufficient to understand the kinetics and thermodynamics of these substates and their role in complexation.

In this regard the current article investigates the conformational heterogeneity in P450cam by considering both ‘closed’ and ‘open’ crystallographic poses of substrate-free P450cam at equal footing. Via spawning unbiased equilibrium MD simulation trajectories from both crystal structures, we explore the possibilities of each crystallographic conformation of making dynamical interconversion to the other crystallographic conformation and build an exhaustive conformational ensemble of substrate-free conformation. A Markovian model based upon cumulative 84 *μs* of atomistic MD data of P450cam quantitatively maps the underlying conformational landscape and identifies multiple key macrostates in the landscape. In particular, the model recovers both ‘closed’ conformation and ‘open’ conformation as major macrostates in the free energy landscape, with the ‘closed’ conformation estimated to be the most populated. The model also predicts a dynamical equilibrium of transition between ‘open’ and ‘closed’ macrostate. However, the rate of *closed* → *open* transition is found to be considerably slower than the reverse (*open* → *closed*) transition. An important finding of current work is that the *direct* transition between open (O) and closed (C) conformation is not the preferred route. Rather, our work predicts that the transition is facilitated via an intermediate (I) macrostate, which is also spatially positioned between these two macrostates. This can also be seen in figure 6, where major contribution to the net flux is going from open (O) to close (C) conformation via intermediate (I). The current work’s prediction of a dynamic equilibrium between closed and open conformation of substrate-free P450cam, with the equilibrium tilted towards closed conformation, suggest an interesting functional implication related to substrate-recognition process in cytochrome P450cam: while substrate like camphor would visit both open and closed conformations of P450cam, it would mostly encounter the closed conformation due to its significantly higher chances of occurrences than other populated macrostates and hence substrate needs to find an opening to the heme via an induced fit based mechanism. However, the presence of dynamic equilibrium between closed and open macrostate also suggests a finite, albeit low, possibility of conformational selection based mechanism for substrate recognition.

### Comparison with prior experimental and computational investigations

The quantification of conformational heterogeneity in the current work reconciles multiple experimental investigations and theoretical works pertaining to P450cam. First, the observation of an intermediate macrostate and a relatively broader distribution of conformational space in open macrostate, are in line with Goodin and co-workers’ previous report of crystallographic structure of ‘open’ conformation of substrate-free P450cam(15) and hypothesis of existence of multiple conformations accessible during substrate recognition(16). In particular, Goodin and coworkers(16) used crystal structures bound to a library of tethered substrates and found that the open conformation is not induced by the tethered substrate but instead dynamically sampled in the substrate-free form. Their principal component analysis of thirty crystal structures also provided the first direct observation of at least three distinct clusters(16) of conformational states closed, intermediate and open and proposed a step-wise closing pathways between them. This observation of multiple conformations is consistent with prediction of current computer simulations. The current results also agree partly with the investigations of P450cam in solution by DEER spectroscopy, which had shown that the substrate-free P450cam exists in the open conformation and binding of substrate leads to a closed conformation (21). Recent NMR studies of substrate-free P450cam(22,23) had revealed the presence of a highly flexible active site which populates an ensemble of conformations based on the segmental movement of F and G helices and suggested presence of a relatively small barrier free energy between these conformations. The IR and UV-visible spectroscopic investigations(24) provide evidence of the conformational heterogeneity in P450cam for both bound and free P450cam and also for the existence of substrate-free P450cam in at least three distinct conformational states the open, the closed and the intermediate state. However, they observed that the conformational ensemble of substrate-free P450cam is highly dominated by the closed conformational state, thereby lending credence to our current work. Together, the present work provides a quantitative perspective to the notion of conformational heterogeneity in substrate-free P450cam, supported by multiple prior spectroscopic and crystallographic investigations. Additionally, the current work elucidates the possible unfolding of B’ helix, predominantly from open conformations, which is in line with earlier experimental reports of B’ helix exhibiting higher thermal motion(14) and being disordered in open form in absence of substrate.(15). More over, comparison of multiple existing cytochrome P450 structures showed the well preserved protein fold but differing in the orientation of B’ helix and its secondary structure state (folded or unfolded) (31). Besides, observation of a disordered B’ helix during substrate-egress in previous computer simulation(33) also provides key support to the current result (using a different protein force field than the current work). Therefore, as quantified in the current work, the dynamics of B’ helix also plays an important role in defining the conformational heterogeneity in cytochrome P450.

We note that previous computational investigations(33,34), based upon biased MD simulations, have suggested diverse insights on conformational plasticity in P450cam and its related implication in substrate recognition processes. Rydzewski and coworkers(33) have explored the egress pathways of substrate from the closed cavity using metadynamics simulation and showed that substrate exit from the heme cavity leads to a transition from a heterogeneous ensemble of closed substrate-bound conformers to the basin comprising of the open conformations of P450cam. Based on the observation of shift of conformation from closed to open state during substrate egress, the work suggested an induced-fit based substrate recognition mechanism. On the other hand, Markwick and coworkers,(34) using adaptive accelerated molecular dynamics simulation approach, have indicated that while closed conformation are free energetically more stable, substrate-free closed protein can potentially access a fully open and partially open conformation and suggested induced-fit and conformational selection based mechanism for binding small and large substrate respectively. Our results contradict their prediction that substrate-free open protein can not access a closed structure at all. On the other hand, the prediction of current work gels well with the ‘induced fit’ mechanism proposed by Ahalawat and co-workers (32) and by Dandekar et. al. (55) of substrate-recognition process by P450cam. However, none of these works has simultaneously considered spawning unbiased trajectories from both closed and open crystallographic poses for exploring their interconversion, as has been done in the current work.

### Relevance to other cytochrome P450 family

Finally, the proposed role of active site flexibility in substrate recognition in P450 family goes beyond P450cam. The substrate specificities across family of P450s show strong diversity and are believed to largely result from differences in structure, flexibility and dynamics of substrate access channel that connects the protein surface to the deeply buried heme centre. Especially, significant diversities have been noted in the sequence of the substrate-binding channel across various forms of P450(57), and as a result, the structure and flexibility of the substrate binding channel varies considerably from one form to another. For example, the small closed substrate access channels seen in many prokaryotic P450s (12, 58–60) are in marked contrast with the bigger and more wide access channels for mammalian microsomal enzymes involved in drug metabolism(61–63). Apart from P450cam, investigations have revealed several examples of bacterial P450s that exist both in an open conformation in the absence of substrate and in a closed conformation in the presence of substrate or ligand(26, 62, 64–69). Hence it is expected that the extent of dynamical heterogeneity that current work has discovered in the conformational ensemble of P450cam (which was long perceived to have narrow substrate channel), is relevant and can be more pronounced in P450s with relatively large access channels. Evidences both supporting and refuting the conformational change associated with substrate recognition processes exist among drug-metabolising microcosmal P450s. For example, while P450 CYP2B4 has been reported to undergo significant conformational change upon substrate-recognition(27, 62, 63), CYP3A4 P450(70) has been observed in both the substrate-bound and -free forms with relatively small changes in structures. Together, these observations point to a greater functional role of conformational heterogeneity in dictating the mechanism of substrate recognition in cytochrome P450.

## AUTHOR CONTRIBUTION

JM, BD and NA designed the project. BD performed the computer simulations. JM, BD and NA analysed the results and wrote the paper.

## SUPPLEMENTAL INFORMATION

Supporting information with figures illustrating adaptive sampling approach implemented in the current work, FES along TICA projection, VAMP-2 score as a function of number of cluster centres, bidirectional rate network among macrostates, snapshot representing important residue interactions stabilising the macrostates

## ACKNOWLEDGEMENTS

This work was supported by computing resources obtained from shared facility of TIFR Center for Interdisciplinary Sciences, India. We acknowledge support of the Department of Atomic Energy, Government of India, under Project Identification No. RTI 4007. JM would like to acknowledge research grants received from Ramanujan Fellowship and core research grant provided by the Department of Science and Technology (DST) of India (CRG/2019/001219).

## REFERENCES

1. Guengerich, F. P., M. R. Waterman, and M. Egli, 2016. Recent Structural Insights into Cytochrome P450 Function. Trends in Pharmacological Sciences 37:625–640. https://doi.org/10.1016/j.tips.2016.05.006.

2. Manikandan, P., and S. Nagini, 2018. Cytochrome P450 Structure, Function and Clinical Significance: A Review. Current Drug Targets 19:38–54. https://doi.org/10.2174/1389450118666170125144557.

3. Coon, M. J., 2005. Cytochrome P450: Nature’s Most Versatile Biological Catalyst. Annu Rev. Pharmaco. 45:1–25. https://doi.org/10.1146/annurev.pharmtox.45.120403.100030.

4. Wade, R. C., D. Motiejunas, K. Schleinkofer Sudarko, P. J. Winn, A. Banerjee, A. Kariakin, and C. Jung, 2005. Multiple molecular recognition mechanisms. Cytochrome P450 - A case study. Biochimica et Biophysica Acta (BBA) - Proteins and Proteomics 1754:239–244. http://www.sciencedirect.com/science/article/pii/S1570963905003080.

5. Miao, Y., Z. Yi, C. Cantrell, D. Glass, J. Baudry, N. Jain, and J. Smith, 2012. Coupled Flexibility Change in Cytochrome P450cam Substrate Binding Determined by Neutron Scattering, NMR, and Molecular Dynamics Simulation. Biophys. J 103:2167–2176. http://www.sciencedirect.com/science/article/pii/S0006349512011174.

6. Miao, Y., and J. Baudry, 2011. Active-Site Hydration and Water Diffusion in Cytochrome P450cam: A Highly Dynamic Process. Biophys. J 101:1493–1503. http://www.sciencedirect.com/science/article/pii/S0006349511009660.

7. Arnold, F. H., 2019. Innovation by Evolution: Bringing New Chemistry to Life (Nobel Lecture). Angewandte Chemie International Edition 58:14420–14426. https://onlinelibrary.wiley.com/doi/abs/10.1002/anie.201907729.

8. Li, A., C. G. Acevedo-Rocha, L. D. Amore, J. Chen, Y. Peng, M. Garcia-Borras, C. Gao, J. Zhu, H. Rickerby, S. Osuna, J. Zhou, and M. T. Reetz, 2020. Regio- and Stereoselective Steroid Hydroxylation at C7 by Cytochrome P450 Monooxygenase Mutants. Angewandte Chemie International Edition 59:12499–12505. https://onlinelibrary.wiley.com/doi/abs/10.1002/anie.202003139.

9. Zhou, H., B. Wang, F. Wang, X. Yu, L. Ma, A. Li, and M. T. Reetz, 2019. Chemo- and Regioselective Dihydroxylation of Benzene to Hydroquinone Enabled by Engineered CytochromeP450 Monooxygenase. Angewandte Chemie International Edition 58:764–768. https://onlinelibrary.wiley.com/doi/abs/10.1002/anie.201812093.

10. Griffin, B. W., and J. A. Peterson, 1972. Camphor binding of Pseudomonas putida cytochrome P-450. Kinetics and thermodynamics of the reaction. Biochemistry 11:4740–4746. https://doi.org/10.1021/bi00775a017, pMID: 4655251.

11. Helms, V., and R. C. Wade, 1995. Thermodynamics of water mediating protein-ligand interactions in cytochrome P450cam: a molecular dynamics study. Biophys. J 69:810–824. http://dx.doi.org/10.1016/S0006-3495(95)79955-6.

12. Poulos, T. L., B. C. Finzel, and A. J. Howard, 1987. High-resolution crystal structure of cytochrome P450cam. J. Mol. Biol 195:687–700. http://www.sciencedirect.com/science/article/pii/0022283687901902.

13. Raag, R., and T. L. Poulos, 1991. Crystal structures of cytochrome p-450cam complexed with camphane,thiocamphor,and adamantane,factors controlling p-450 substrate hydroxylation. Biochemistry 30:2674–2684.

14. Poulos, T. L., B. C. Finzel, and A. J. Howard, 1986. Crystal structure of substrate-free Pseudomonas putida cytochrome P-450. Biochemistry 25:5314–5322. https://doi.org/10.1021/bi00366a049, pMID: 3768350.

15. Lee, Y.-T., R. F. Wilson, I. Rupniewski, and D. B. Goodin, 2010. P450cam Visits an Open Conformation in the Absence of Substrate,. Biochemistry 49:3412–3419. https://doi.org/10.1021/bi100183g, pMID: 20297780.

16. Lee, Y.-T., E. C. Glazer, R. F. Wilson, C. D. Stout, and D. B. Goodin, 2011. Three Clusters of Conformational States in P450cam Reveal a Multistep Pathway for Closing of the Substrate Access Channel,. Biochemistry 50:693–703. https://doi.org/10.1021/bi101726d, pMID: 21171581.

17. Groves, J. T., 1985. Key elements of the chemistry of cytochrome-P-450: The oxygen rebound mechanism. J. Chem. Educ 62:928–931.

18. Schlichting, I., J. Berendzen, K. Chu, A. M. Stock, S. A. Maves, D. E. Benson, R. M. Sweet, D. Ringe, G. A. Petsko, and S. G. Sligar, 2000. The Catalytic Pathway of Cytochrome P450cam at Atomic Resolution. Science 287:1615–1622. https://science.sciencemag.org/content/287/5458/1615.

19. Skinner, S. P., W.-M. Liu, Y. Hiruma, M. Timmer, A. Blok, M. A. S. Hass, and M. Ubbink, 2015. Delicate conformational balance of the redox enzyme cytochrome P450cam. Proc. Natl. Acad. Sci. U.S.A 112:9022–9027. https://www.pnas.org/content/112/29/9022.

20. Dunn, A. R., I. J. Dmochowski, A. M. Bilwes, H. B. Gray, and B. R. Crane, 2001. Probing the open state of cytochrome P450cam with ruthenium-linker substrates. Proc. Natl. Acad. Sci. U.S.A 98:12420–12425. https://www.pnas.org/content/98/22/12420.

21. Stoll, S., Y. T. Lee, M. Zhang, R. F. Wilson, R. D. Britt, and D. B. Goodin, 2012. Double electron-electron resonance shows cytochrome P450cam undergoes a conformational change in solution upon binding substrate. Proc. Natl. Acad. Sci. U.S.A 109:12888–12893.

22. Asciutto, E. K., M. J. Young, J. Madura, S. S. Pochapsky, and T. C. Pochapsky, 2012. Solution structural ensembles of substrate-free cytochrome P450 cam. Biochemistry 51:3383–3393.

23. Colthart, A. M., D. R. Tietz, Y. Ni, J. L. Friedman, M. Dang, and T. C. Pochapsky, 2016. Detection of substrate-dependent conformational changes in the P450 fold by nuclear magnetic resonance. Sci. Rep. 6:1–11. http://dx.doi.org/10.1038/srep22035.

24. Basom, E. J., B. A. Manifold, and M. C. Thielges, 2017. Conformational Heterogeneity and the Affinity of Substrate Molecular Recognition by Cytochrome P450cam. Biochemistry 56:3248–3256.

25. Savino, C., L. C. Montemiglio, G. Sciara, A. E. Miele, S. G. Kendrew, P. Jemth, S. Gianni, and B. Vallone, 2009. Investigating the Structural Plasticity of a Cytochrome P450: Three-Dimentional Structures of P450 EryK and Binding to its Physiological Substrates. J. Biol. Chem. 284:29170–29179.

26. Sherman, D. H., S. Li, L. V. Yermalitskaya, Y. Kim, J. A. Smith, M. R. Waterman, and L. M. Podust, 2006. The Structural Basis for Substrate Anchoring, Active Site Selectivity, and Product Formation by P450 PikC from Streptomyces venezuelae. J. Biol. Chem. 281:26289–26297. http://www.jbc.org/content/281/36/26289.abstract.

27. Zhao, Y., M. A. White, B. K. Muralidhara, L. Sun, J. R. Halpert, and C. D. Stout, 2006. Structure of Microsomal Cytochrome P450 2B4 Complexed with the Antifungal Drug Bifonazole: Insights into P450 Conformational Plasticity and Membrane Interaction. J. Biol. Chem. 281:5973–5981. http://www.jbc.org/content/281/9/5973.abstract.

28. Husic, B. E., and V. S. Pande, 2018. Markov State Models: From an Art to a Science. J. Am. Chem. Soc. 140:2386–2396. https://doi.org/10.1021/jacs.7b12191,pMID:29323881.

29. Bowman, G. R., V. S. Pande, and F. Noé, 2014. An Introduction to Markov State Models and Their Application to Long Timescale Molecular Simulation. In Adv. Exp. Med. Biol.

30. Chodera, J. D., and F. Noé, 2014. Markov state models of biomolecular conformational dynamics. Curr. Opin. Struc. Biol. 25:135–144. https://doi.org/10.1016%2Fj.sbi.2014.04.002.

31. de Montellano, P. R. O., editor, 2015. Cytochrome P450: Structure, Mechanism and Biochemistry. Springer International Publishing. https://doi.org/10.1007/978-3-319-12108-6.

32. Ahalawat, N., and J. Mondal, 2018. Mapping the Substrate Recognition Pathway in Cytochrome P450. J. Am. Chem. Soc. 140:17743–17752. https://doi.org/10.1021/jacs.8b10840.

33. Rydzewski, J., and W. Nowak, 2017. Thermodynamics of camphor migration in cytochrome P450cam by atomistic simulations. Sci. Rep. 7:7736.

34. Markwick, P. R. L., L. C. T. Pierce, D. B. Goodin, and J. A. McCammon, 2011. Adaptive Accelerated Molecular Dynamics (Ad-AMD) Revealing the Molecular Plasticity of P450cam. J. Phys. Chem. Lett 2:158–164. https://doi.org/10.1021/jz101462n, pMID: 21307966.

35. Sali, A., and T. L. Blundell, 1993. Comparative Protein Modelling by Satisfaction of Spatial Restraints. J. Mol. Biol 234:779–815. http://www.sciencedirect.com/science/article/pii/S0022283683716268.

36. Mackerell, A. D., M. Feig, and C. L. Brooks, 2004. Extending the treatment of backbone energetics in protein force fields: Limitations of gas-phase quantum mechanics in reproducing protein conformational distributions in molecular dynamics simulations. J. Comput. Chem 25:1400–1415. https://doi.org/10.1002/jcc.20065.

37. Jorgensen, W. L., J. Chandrasekhar, J. D. Madura, R. W. Impey, and M. L. Klein, 1983. Comparison of simple potential functions for simulating liquid water. J. Chem. Phys. 79:926–935.

38. Pail, S., and B. Hess, 2013. A flexible algorithm for calculating pair interactions on {SIMD} architectures. Comput. Phys. Comm.s 184:2641–2650. http://www.sciencedirect.com/science/article/pii/S0010465513001975.

39. Darden, T., D. York, and L. G. Pederson, 1993. Particle mesh Ewald: An N?log(N) method for Ewald sums in large systems. J. Chem. Phys. 98:952.

40. Miyamoto, S., and P. Kollman, 1992. Settle: An analytical version of the SHAKE and RATTLE algorithm for rigid water models. J. Comput. Chem. 13:952–962.

41. Bussi, G., D. Donadio, and M. Parrinello, 2007. Canonical sampling through velocity rescaling. The Journal of Chemical Physics 126:014101. https://doi.org/10.1063/1.2408420.

42. Parrinello, M., and A. Rahman, 1981. Polymorphic transitions in single crystals: A new molecular dynamics method. J. Appl. Phys 52:7182–7190. http://scitation.aip.org/content/aip/journal/jap/52/12/10.1063/1.328693.

43. Abraham, M. J., T. Murtola, R. Schulz, S. Pall, J. C. Smith, B. Hess, and E. Lindahl, 2015. GROMACS: High performance molecular simulations through multi-level parallelism from laptops to supercomputers. SoftwareX 1-2:19–25. http://www.sciencedirect.com/science/article/pii/S2352711015000059.

44. Scherer, M. K., B. Trendelkamp-Schroer, F. Paul, G. Perez-Hernandez, M. Hoffmann, N. Plattner, C. Wehmeyer, J.-H. Prinz, and F. Noe, 2015. PyEMMA 2: A Software Package for Estimation, Validation, and Analysis of Markov Models. J. Chem. Theory Comput 11:5525–5542. http://dx.doi.org/10.1021/acs.jctc.5b00743.

45. Wu, H., and F. Noé, 2019. Variational Approach for Learning Markov Processes from Time Series Data. Journal of Nonlinear Science 30:23–66. https://doi.org/10.1007/s00332-019-09567-y.

46. Paul, F., H. Wu, M. Vossel, B. L. de Groot, and F. Noé, 2019. Identification of kinetic order parameters for non-equilibrium dynamics. The Journal of Chemical Physics 150:164120. https://doi.org/10.1063/1.5083627.

47. Lloyd, S., 2006. Least Squares Quantization in PCM. IEEE Trans. Inf. Theor. 28:129–137. http://dx.doi.org/10.1109/TIT.1982.1056489.

48. Röblitz, S., and M. Weber, 2013. Fuzzy spectral clustering by PCCA+: application to Markov state models and data classification. Adv. Data Anal. Classi. 7:147–179. https://doi.org/10.1007/s11634-013-0134-6.

49. Polizzi, N. F., M. J. Therien, and D. N. Beratan, 2016. Mean First-Passage Times in Biology. Israel Journal of Chemistry 56:816–824.

50. Singhal, N., and V. S. Pande, 2005. Error analysis and efficient sampling in Markovian state models for molecular dynamics. J. Chem. Phys. 123:204909. https://doi.org/10.1063/1.2116947.

51. E., W., and E. Vanden-Eijnden, 2006. Towards a Theory of Transition Paths. Journal of Statistical Physics 123:503. https://doi.org/10.1007/s10955-005-9003-9.

52. Metzner, P., C. Schutte, and E. Vanden-Eijnden, 2009. Transition Path Theory for Markov Jump Processes. Multiscale Modeling & Simulation 7:1192–1219.

53. Noé, F., C. Schütte, E. Vanden-Eijnden, L. Reich, and T. R. Weikl, 2009. Constructing the equilibrium ensemble of folding pathways from short off-equilibrium simulations. Proc. Natl. Acad. Sci. U.S.A 106:19011–19016. http://www.pnas.org/content/106/45/19011.

54. Wehmeyer, C., M. K. Scherer, T. Hempel, B. E. Husic, S. Olsson, and F. Noe, 2018. Introduction to Markov state modeling with the PyEMMA software [Article v1.0]. Living Journal of Computational Molecular Science 1:5965.

55. Dandekar, B. R., and J. Mondal, 2020. Capturing Protein–Ligand Recognition Pathways in Coarse-Grained Simulation. The Journal of Physical Chemistry Letters 11:5302–5311. https://doi.org/10.1021/acs.jpclett.0c01683, pMID: 32520567.

56. Guengerich, F. P., S. A. Child, I. R. Barckhausen, and M. H. Goldfarb, 0. Kinetic Evidence for an Induced-Fit Mechanism in the Binding of the Substrate Camphor by Cytochrome P450cam. ACS Catalysis 0:639–649. https://doi.org/10.1021/acscatal.0c04455.

57. Gotoh, O., 1992. Substrate recognition sites in cytochrome P450 family 2 (CYP2) proteins inferred from comparative analyses of amino acid and coding nucleotide sequences. J. Biol. Chem. 267:83–90. http://www.jbc.org/content/267/1/83.abstract.

58. Meharenna, Y. T., H. Li, D. B. Hawkes, A. G. Pearson, J. De Voss, and T. L. Poulos, 2004. Crystal Structure of P450cin in a Complex with Its Substrate, 1,8-Cineole, a Close Structural Homologue to d-Camphor, the Substrate for P450cam,. Biochemistry 43:9487–9494. https://doi.org/10.1021/bi049293p, pMID: 15260491.

59. Pylypenko, O., and I. Schlichting, 2004. Structural Aspects of Ligand Binding to and Electron Transfer in Bacterial and Fungal P450s. Annu. Rev. Biochem 73:991–1018. https://doi.org/10.1146/annurev.biochem.73.011303.073711, pMID: 15189165.

60. Poulos, T. L., J. Cupp-Vickery, and H. Li, 1995. Structural Studies on Prokaryotic Cytochromes P450, Springer US, Boston, MA, 125–150. https://doi.org/10.1007/978-1-4757-2391-5_4.

61. Johnson, E. F., and C. D. Stout, 2005. Structural diversity of human xenobiotic-metabolizing cytochrome P450 monooxygenases. Biochem. Biophys. Res. Commun 338:331–336. http://www.sciencedirect.com/science/article/pii/S0006291X05019169, celebrating 50 Years of Oxygenases.

62. Scott, E. E., Y. A. He, M. R. Wester, M. A. White, C. C. Chin, J. R. Halpert, E. F. Johnson, and C. D. Stout, 2003. An open conformation of mammalian cytochrome P450 2B4 at 1.6-Å resolution. Proc. Natl. Acad. Sci. U.S.A 100:13196–13201. https://www.pnas.org/content/100/23/13196.

63. Scott, E. E., M. A. White, Y. A. He, E. F. Johnson, C. D. Stout, and J. R. Halpert, 2004. Structure of Mammalian Cytochrome P450 2B4 Complexed with 4-(4-Chlorophenyl)imidazole at 1.9 angstrom Resolution: Insight into the range of P450 conformations and the coordination of redox partner binding. J. Biol. Chem. 279:27294–27301.

64. Li, H., and T. Poulos, 1997. The structure of the cytochrome p450BM-3 haem domain complexed with the fatty acid substrate, palmitoleic acid. Nat. Struct. Mol. Biol. 4:140–146.

65. Zhao, B., F. P. Guengerich, A. Bellamine, D. C. Lamb, M. Izumikawa, L. Lei, L. M. Podust, M. Sundaramoorthy, J. A. Kalaitzis, L. M. Reddy, S. L. Kelly, B. S. Moore, D. Stec, M. Voehler, J. R. Falck, T. Shimada, and M. R. Waterman, 2005. Binding of Two Flaviolin Substrate Molecules, Oxidative Coupling, and Crystal Structure of Streptomyces coelicolor A3(2) Cytochrome P450 158A2. J. Biol. Chem. 280:11599–11607. http://www.jbc.org/content/280/12/11599.abstract.

66. Park, S.-Y., K. Yamane, S. ichi Adachi, Y. Shiro, K. E. Weiss, S. A. Maves, and S. G. Sligar, 2002. Thermophilic cytochrome P450 (CYP119) from Sulfolobus solfataricus: high resolution structure and functional properties. J. Inorg. Biochem 91:491–501. http://www.sciencedirect.com/science/article/pii/S0162013402004464.

67. Xu, L.-H., S. Fushinobu, H. Ikeda, T. Wakagi, and H. Shoun, 2009. Crystal Structures of Cytochrome P450 105P1 from Streptomyces avermitilis: Conformational Flexibility and Histidine Ligation State. Journal of Bacteriology 191:1211–1219. https://jb.asm.org/content/191/4/1211.

68. Ouellet, H., L. M. Podust, and P. R. O. de Montellano, 2008. Mycobacterium tuberculosis CYP130: Crystal Structure, Biophysical Characterisation, and Interactions With Antifungal AZOLE Drugs. J. Biol. Chem. 283:5069–5080. http://www.jbc.org/content/283/8/5069.abstract.

69. Zerbe, K., O. Pylypenko, F. Vitali, W. Zhang, S. Rouset, M. Heck, J. W. Vrijbloed, D. Bischoff, B. Bister, R. D. Sussmuth, S. Pelzer, W. Wohlleben, J. A. Robinson, and I. Schlichting, 2002. Crystal Structure of OxyB, a Cytochrome P450 Implicated in an Oxidative Phenol Coupling Reaction during Vancomycin Biosynthesis. J. Biol. Chem. 277:47476–47485. http://www.jbc.org/content/277/49/47476.abstract.

70. Williams, P. A., J. Cosme, D. M. Vinković, A. Ward, H. C. Angove, P. J. Day, C. Vonrhein, I. J. Tickle, and H. Jhoti, 2004. Crystal Structures of Human Cytochrome P450 3A4 Bound to Metyrapone and Progesterone. Science 305:683–686. https://science.sciencemag.org/content/305/5684/683.

