## Supplemental files for "Reconciling Conformational Heterogeneity and Substrate Recognition in Cytochrome P450"

### Supporting information for "Reconciling Conformational Heterogeneity and Substrate-recognition in Cytochrome P450"

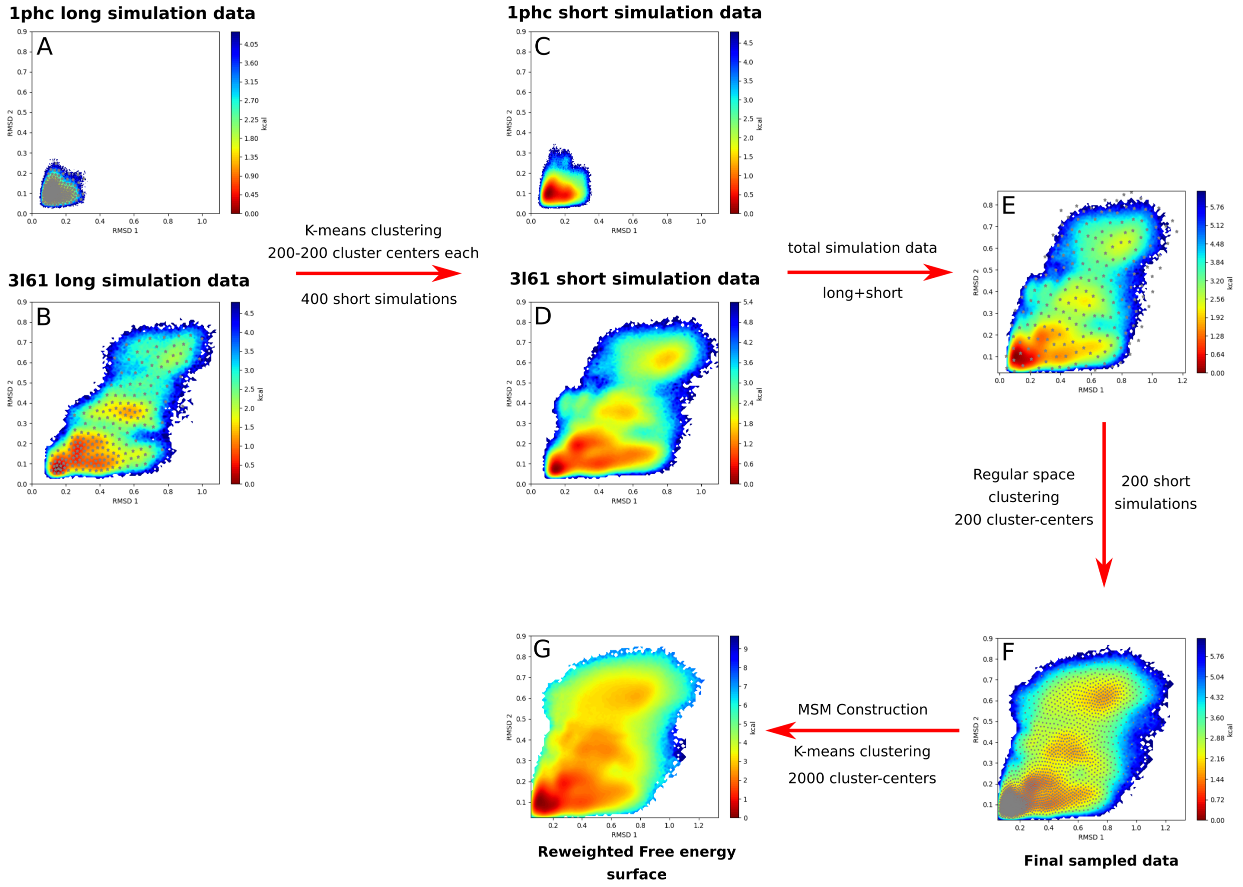


**Figure S1:** Adaptive sampling approach employed to capture conformationally heterogenous free energy surface of cytochrome P450.

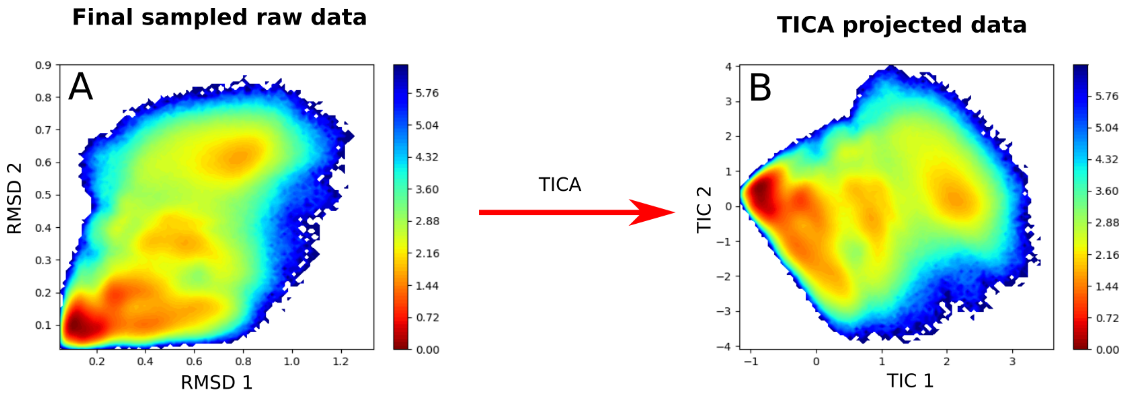


**Figure S2:** The projection of raw data along RMSD1 and RMSD2 onto the linearly transformed TICA dimension TIC1 and TIC2.

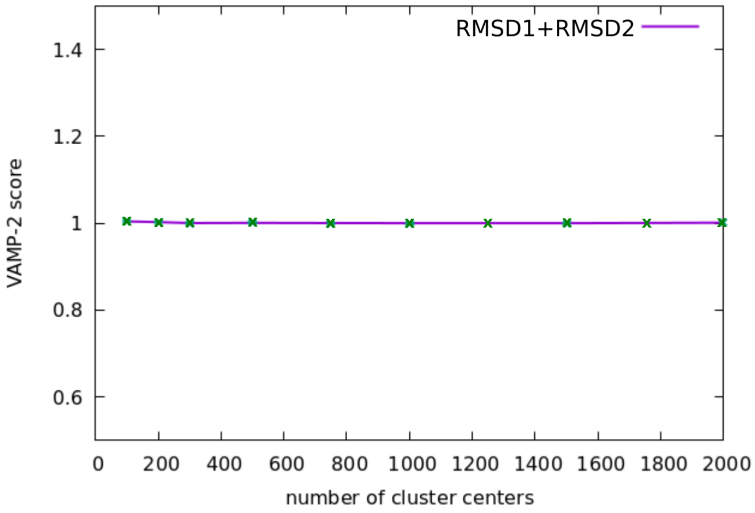


**Figure S3:** The variation of VAMP-2 score as a function of number of cluster-centers.

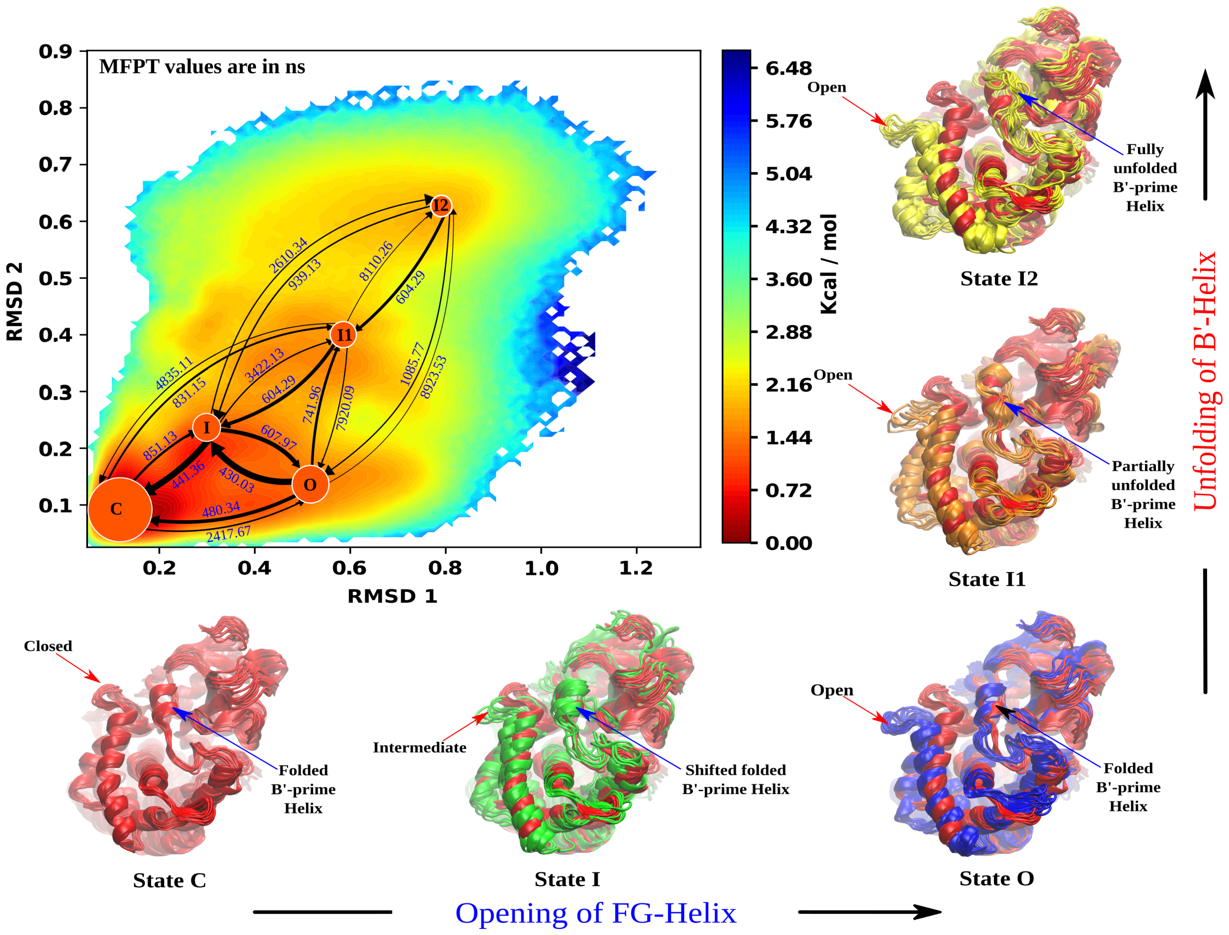


**Figure S4:** Rate network among macrostates: The rate network obtained from MFPT analysis. The network connects five macrostates, the state C (closed), I (intermediate), O (open), I1, I2 and are represented as a circular discs on their respective coordinates (RMSD1,RMSD2). The sizes of the discs are proportional to the stationary population of the respective macrostate. The thickness of the arrows is proportional to the rate (i.e 1/MFPT) of the transition within the macrostates and the MFPT values of corresponding transitions are written over the arrow in ns. The figure also shows the distinct five representative snapshots corresponding to most probable states from each macrostates. The snapshots are shown in comparison with the first state, the closed state C (red colored) to indicate the relative differences in the conformations. In each states 10 snapshots are shown for simplicity.

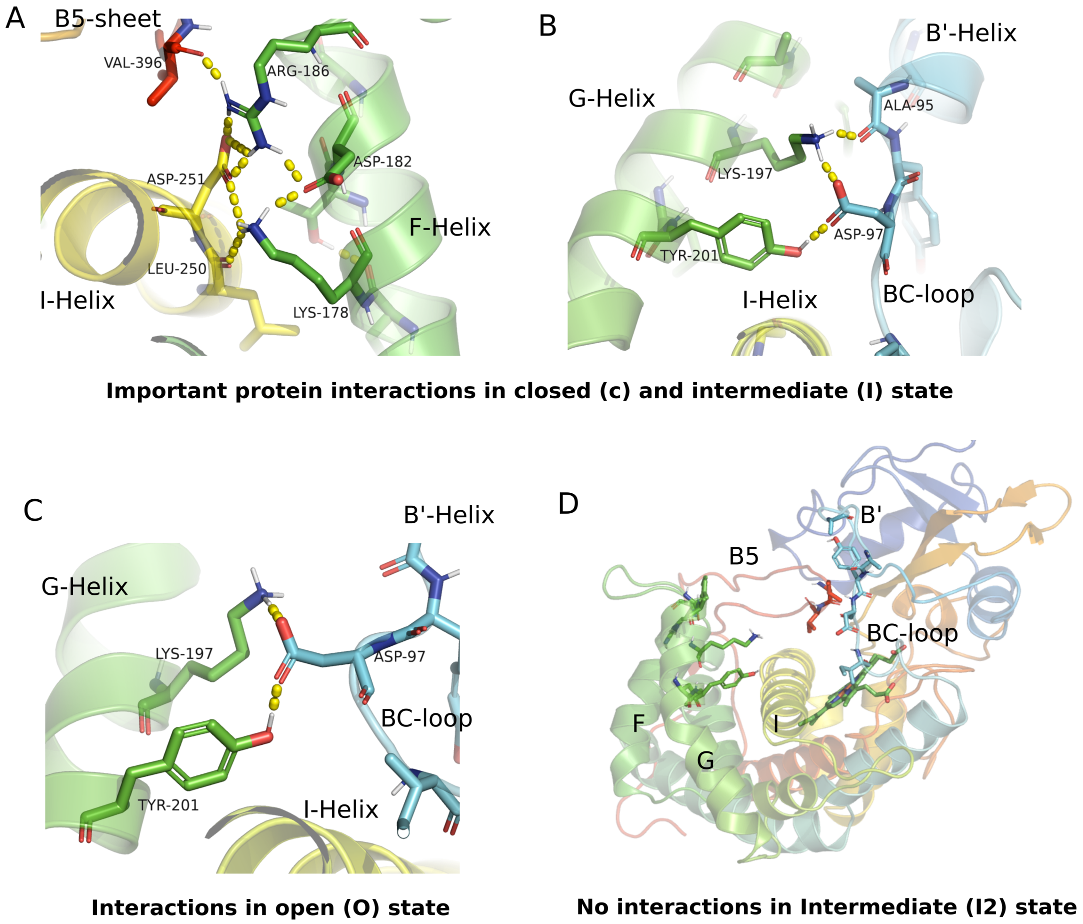


**Figure S5:** Important residue interactions present across the MSM-derived macro states from closed to open states. (A) and (B) Interactions shown in are present in both close(C) and intermediate(I) state. (C) Interactions present in open state. (D) There are loss of above key interactions in partially (I1) and fully open (I2) conformations.
